# Simulating Colloid Motion in Spinning Suspension: Internal Dynamics in Orbital Shaker

**DOI:** 10.1101/2024.02.19.581033

**Authors:** Neil Zhao, Diana Zheng

## Abstract

Colloid suspensions in the form of mammalian or bacterial cell mixtures in orbital shakers are commonly encountered in biomedical research. An understanding of particle motion in these conditions would provide insight into bulk colloid behavior and distribution under short and long timeframes. Such data can be used to supplement and clarify existing concepts of biological phenomena encountered in the laboratory setting. It can also aid biomedical researchers in experiment design and data interpretation. We present a MATLAB based simulation of colloid motion under rotary agitation. Our simulation setup is modular and therefore designed to act as a scaffold that can be customized to simulate different mammalian, bacterial, or particle properties in liquid suspension.

## 1 Introduction

The orbital shaker is commonly used in biological or medical science research as a tool for agitating mixtures into a state of homogeneity [1,2]. Container shape and size have been found to affect mammalian cell growth under rotary mixing [3,4], however there exists a gap in knowledge on the internal dynamics of colloid mixtures in the biological sciences under orbital agitation. Mammalian cells and bacteria are frequently put into liquid suspensions that are transferred to orbital shakers as a step in many standard operating procedures. A common assumption is that an equilibrium of uniform particle distribution is quickly reached. And therefore differences in the local environment around individual cells are negligible in terms of experimental measurements. The error around this assumption naturally increases as the container size increases, the timeframe is decreased, or as the cells start to exhibit behavior that magnify small differences into large outcomes. Cells located closer to the surface might thus be more affected by oxygen diffusion in larger containers. Alternatively, a cell-cell interaction might exist to result in a non-uniform cell density with increasing time.

We present in this manuscript an open-source, MATLAB based simulation of colloid movement under orbital agitation. A basic scaffold of rotary motion, Brownian motion, and particle-particle interaction was constructed. Special cases of conditions found with common mammalian or bacterial cell behavior were then run through the simulation and their results provided. We put forward our simulation as a general platform for researchers to adjust or supplement in modeling colloid motion in spinning suspensions under conditions unique to their own experiments.

## 2 Simulation setup, execution, and results

### 2.1 Mathematical premise

Colloid location at time zero was distributed uniformly over a circle using a random number generator to determine a set of polar coordinates (*r*, θ)_*i*_. The radial coordinates were drawn with a linearly increasing probability proportional to distance from origin. This was done to ensure that the final distribution of colloids would be uniform over a circle.

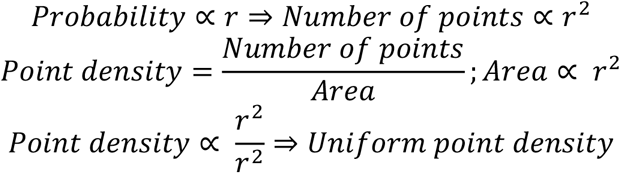

All colloids were given an angular velocity ω around the origin to simulate the overall motion imparted by the orbital shaker.

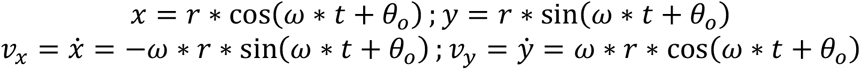

Linear approximation and discretization of the above equations resulted in formats suitable for computer simulation.

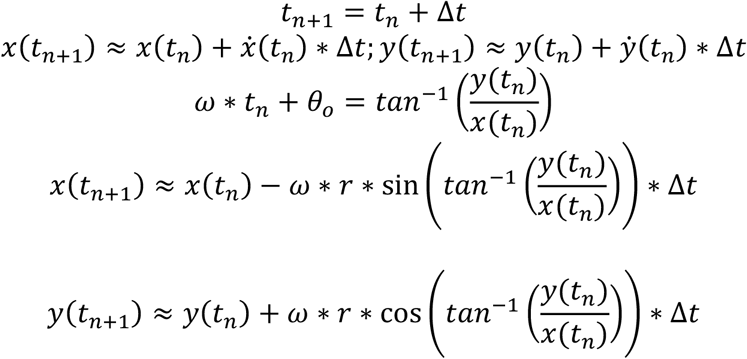

Colloids were also allowed to undergo a random walk at every time step to simulate Brownian motion.

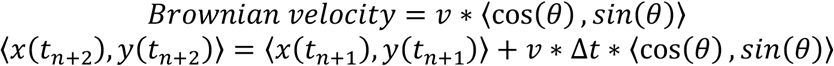

Distance between all colloid combinations was calculated at each step to determine which were within range for adhesion. This was performed by horizontally and vertically concatenating all *x* and *y* coordinates into square matrices. All calculations were then performed as below.

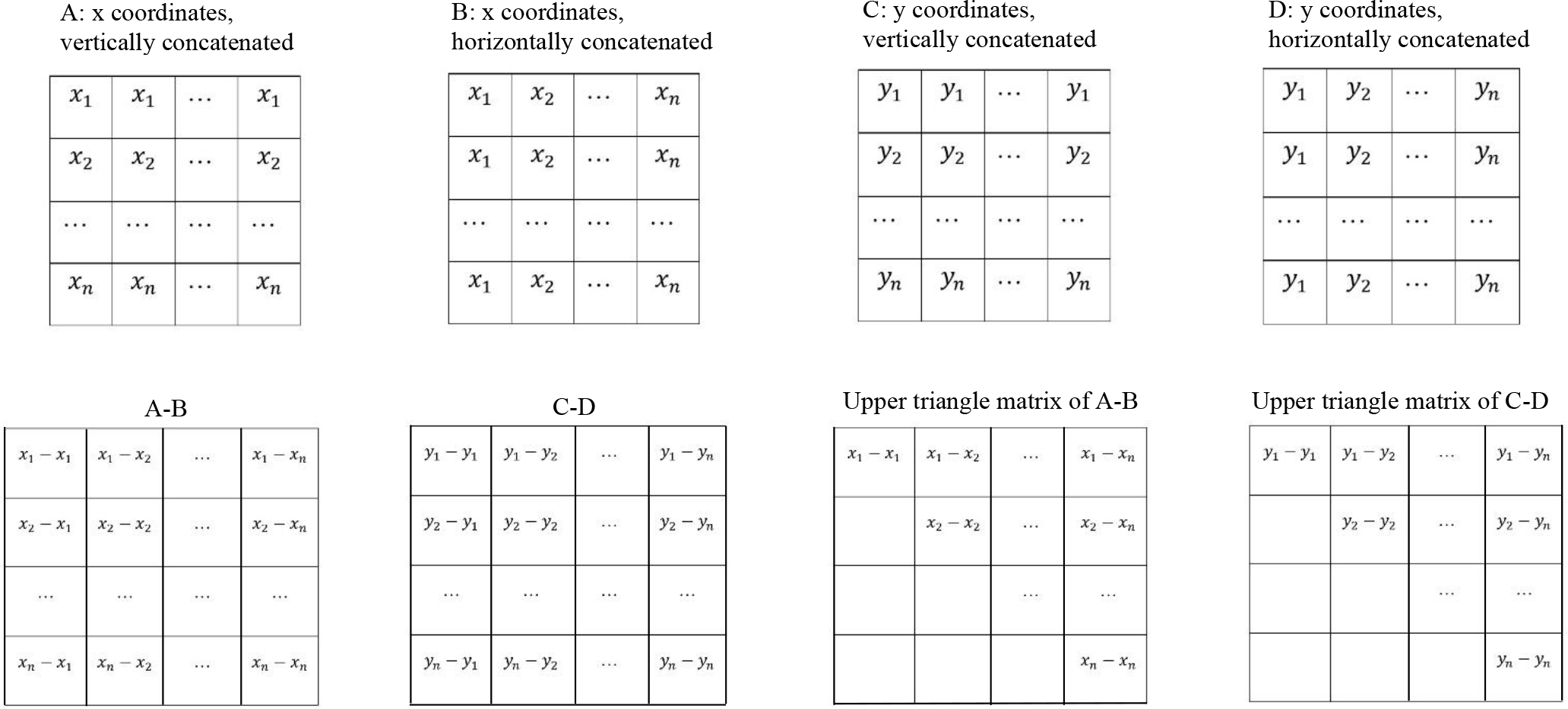

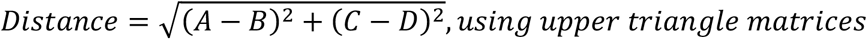

All MATLAB code is provided below with annotations.

### 2.2 MATLAB simulation

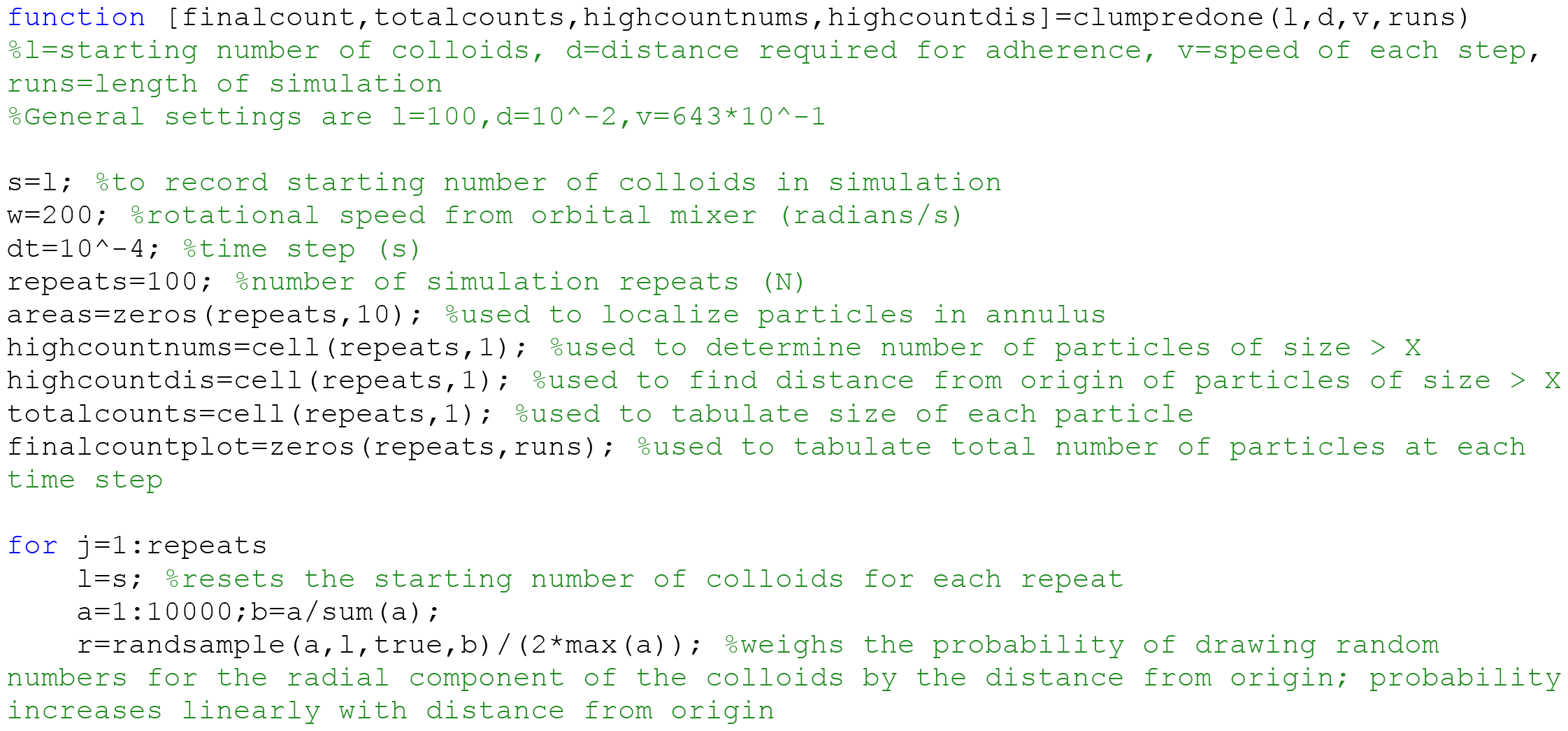
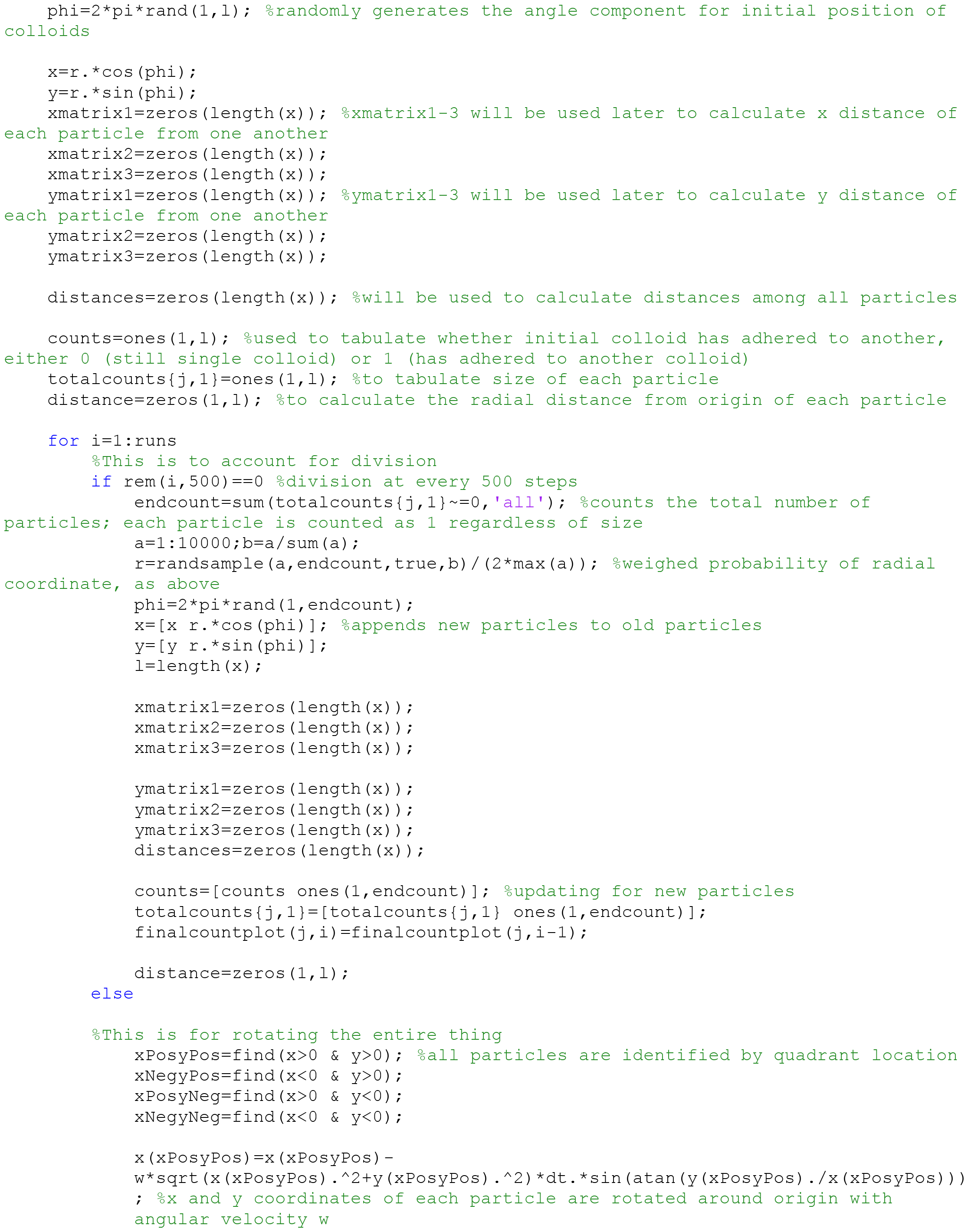
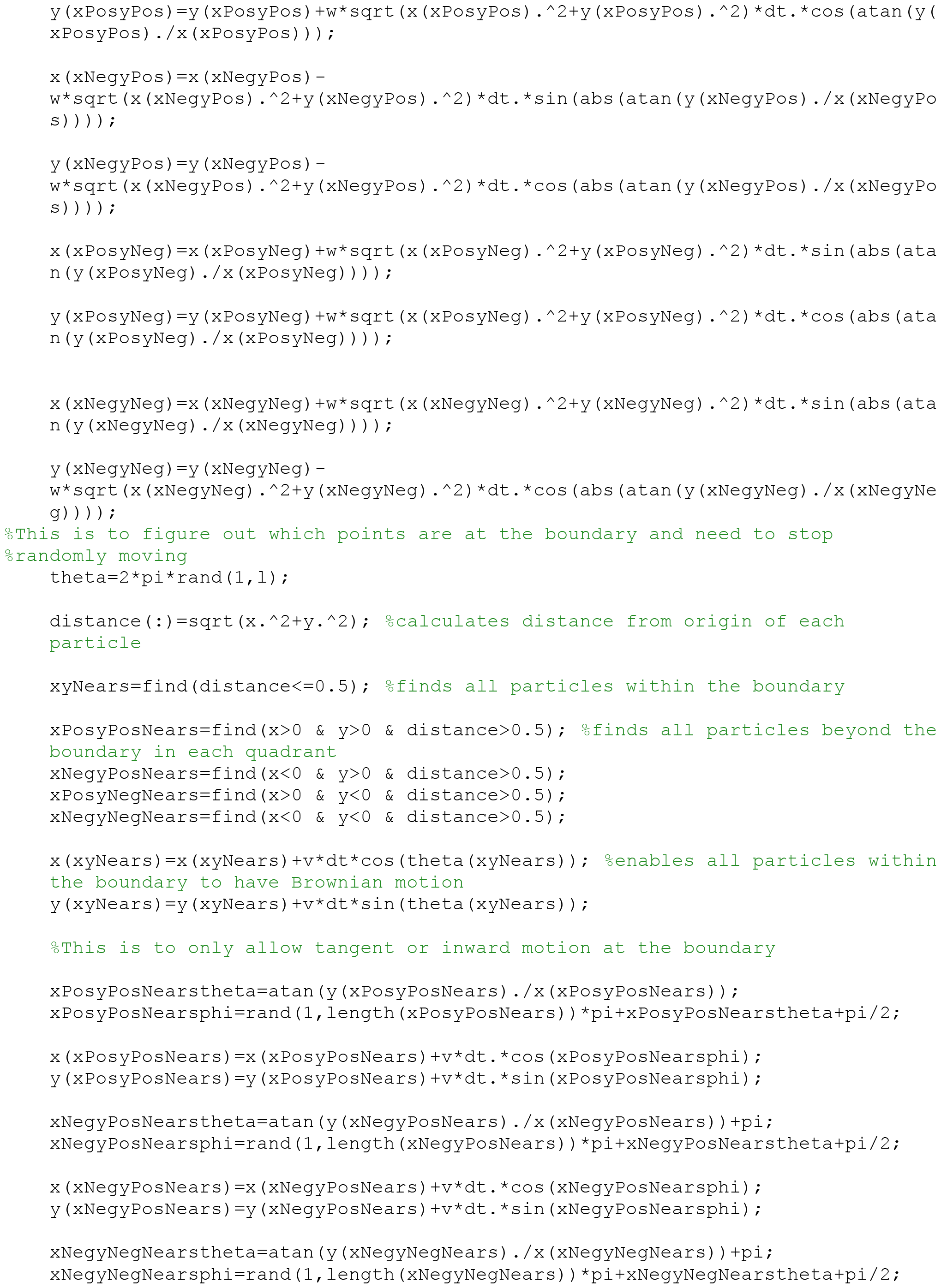
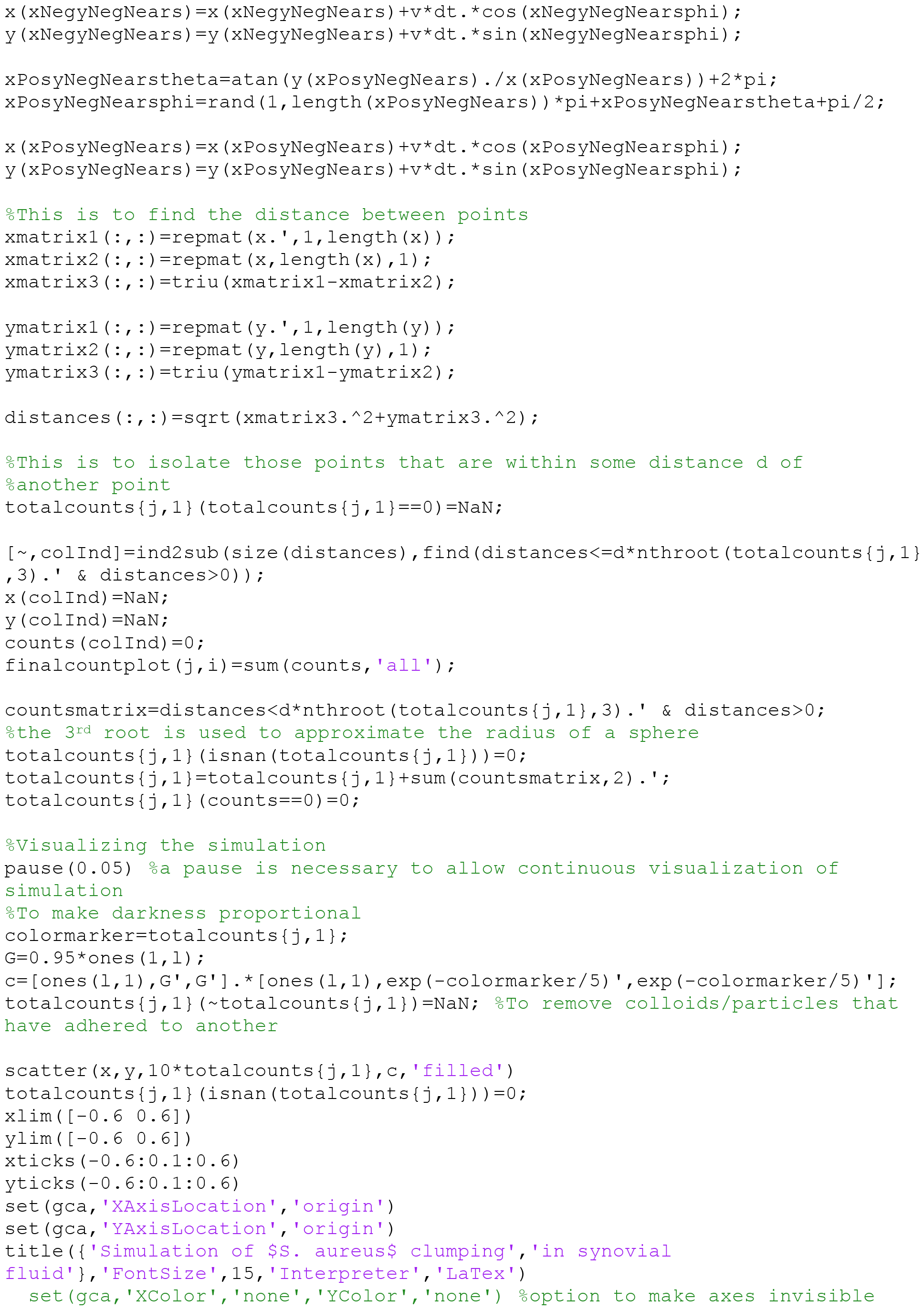
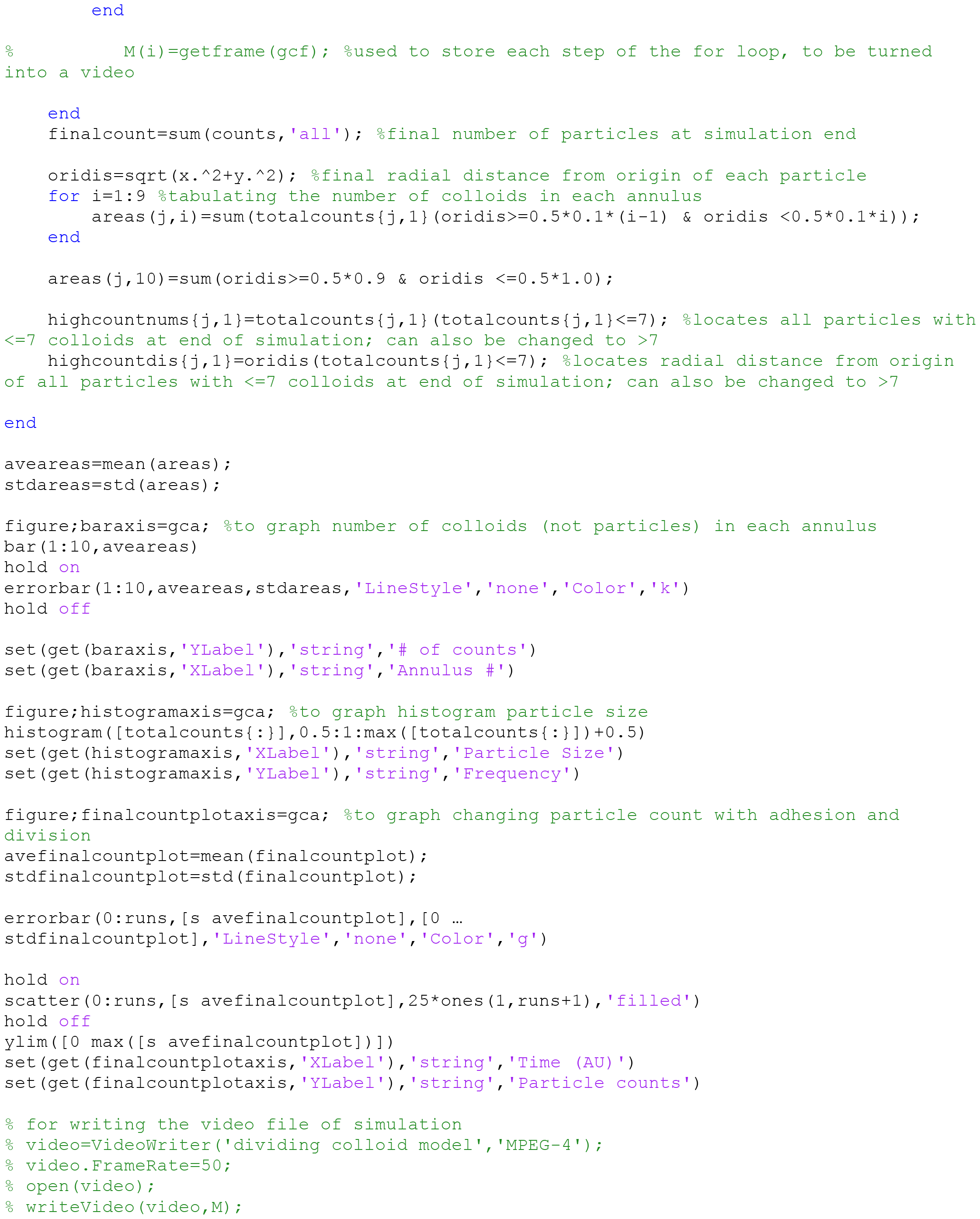
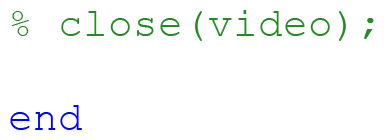

### 2.3 Simulation results

#### Case 1: Colloids in spinning suspension

Colloids were uniformly distributed over a circular area representing common laboratory containers and given Brownian motion and an angular velocity counterclockwise around the origin. Distribution of colloids at the start (Fig. 1A) and end (Fig. 1B) of the simulation showed similar density. Change in colloid position with respect to the origin was measured based on an annular overlay representing increasing radial distances (Fig. 1C). The number of colloids falling into each annulus at the start (Fig. 1D), 1/4 completion (E), ½ completion (F), and end (G) of simulation exhibited a linear increase with radial distance. The area of an annulus increases proportionally with the square of the radial distance. The uniform distribution of colloids resulted in a corresponding increase in colloid numbers proportional to the square of the radial distance, *N* ∝ *r*^2^. The first order Taylor expansion of a parabolic function is linear, which corresponds to our measured change in colloid number with annulus distance.

**Figure 1:**
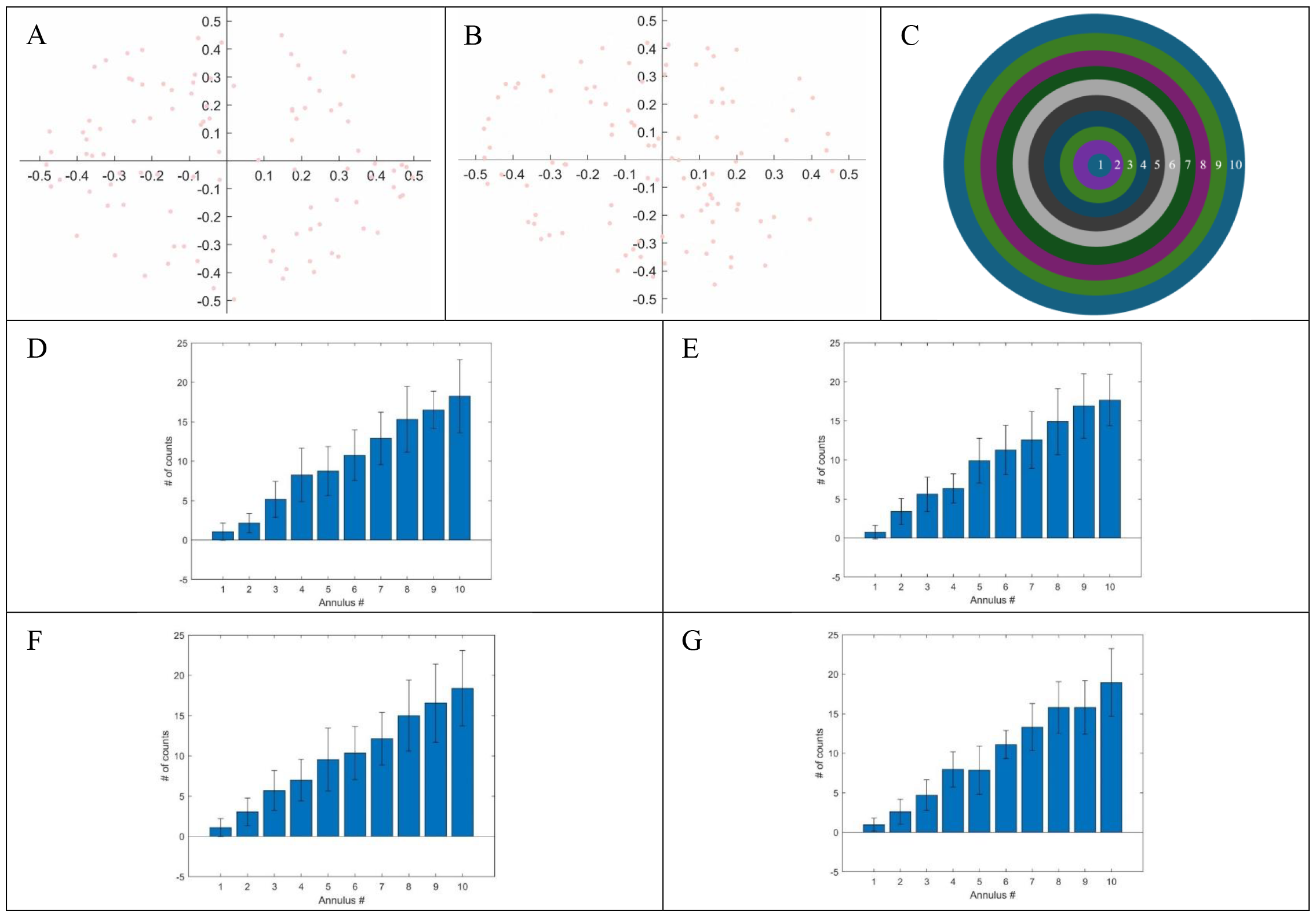
Colloids in a spinning suspension. Distribution of one hundred colloids at the (A) start and (B) end of the simulation. (C) Annulus overlay used to localize colloid position. Position of colloids at the (D) start, (E) 1/4 completion, (F) 1/2 completion, (G) end of the simulation. Plotted are means ± standard deviations. N=20 for all measurements.

#### Case 2: Adhesive colloids in spinning suspension

Colloids with the ability to adhere if the inter-particle distance was less than a defined limit were uniformly distributed over a circular area and allowed to proceed through the simulation as above. We defined particle as the general term for colloids, whether singular or adhered with others. The number of particles decreased throughout the simulation (Fig. 2A-D). Particles formed by the adhesion of multiple colloids were radially distributed about the origin with no apparent angular bias at simulation end. Particle number decreased in a non-linear fashion with respect to time (Fig. 2E). The cumulative particle size distribution of 20 runs exhibited a skewed right, unimodal distribution with a peak at 3 particles (Fig. 2F). A right tail was present, showing <5 particles with >10 colloids at simulation end. Colloid number increased linearly with radial distance from origin (Fig. 3A). Colloid density remained uniform for all annuli, the greatest variation appearing in the two closest annuli to the origin as indicated by the large error bars (Fig. 3B). Small sized particles (colloid number ≤7) and large sized particles (colloid number >7) were both uniformly distributed at simulation end (Fig. 3C,D). Both increased linearly with radial distance from origin, resembling the uniform colloid distribution at simulation art (Fig. 1D).

**Figure 2:**
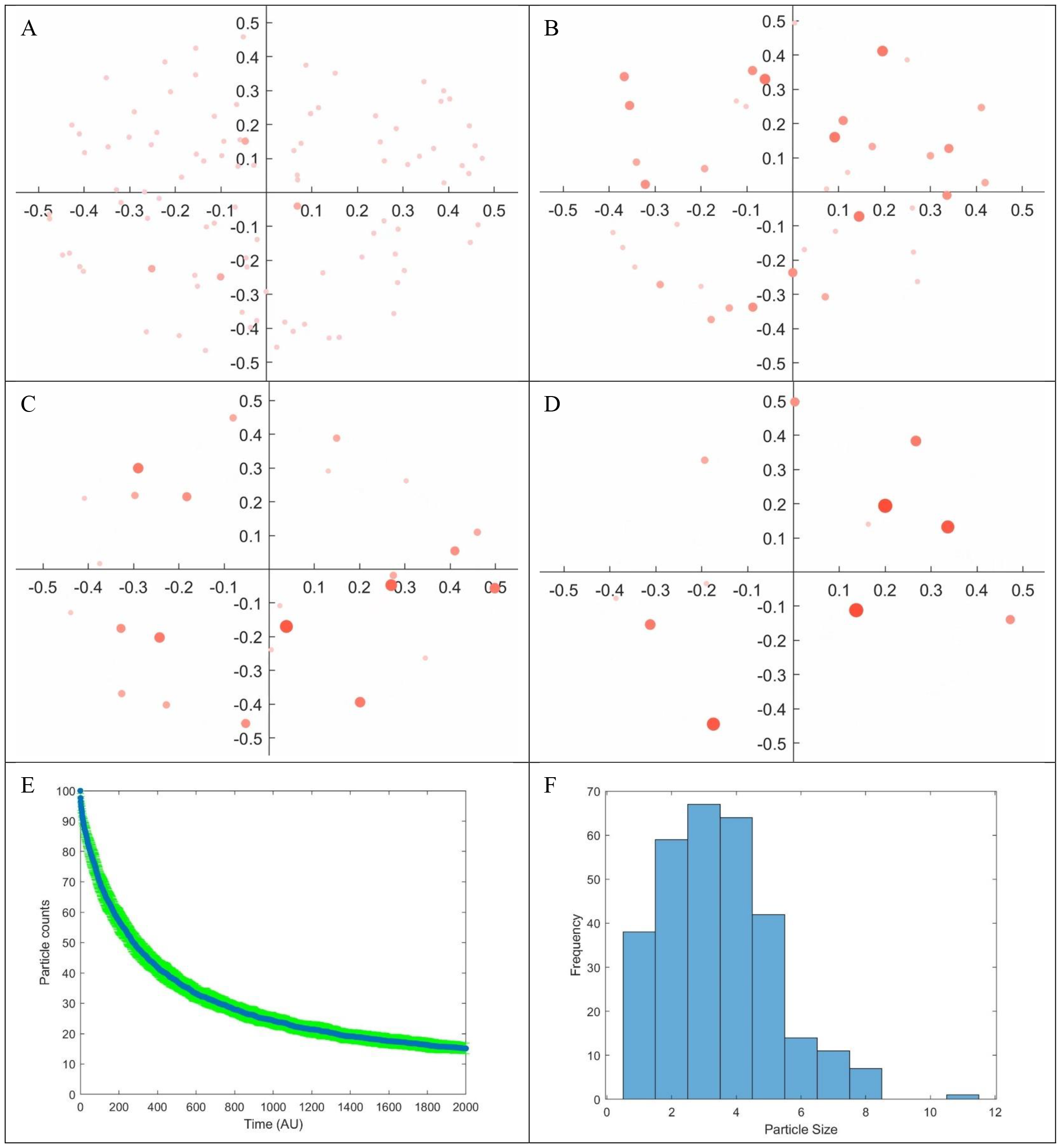
Adhesive colloids in a spinning suspension. Distribution of one hundred adhesive colloids at the (A) start, (B) 1/4 completion, (C) 1/2 completion, (D) end of simulation. Size and color intensity correlate with number of adhered colloids. (E) Change in particle count of adhesive colloids in spinning suspension. Plotted are means ± standard deviations. N=20 for each data point. (F) Distribution of particle size at end of colloid adhesion in spinning suspension simulation. Plotted is the sum of N=20 simulation runs.

**Figure 3:**
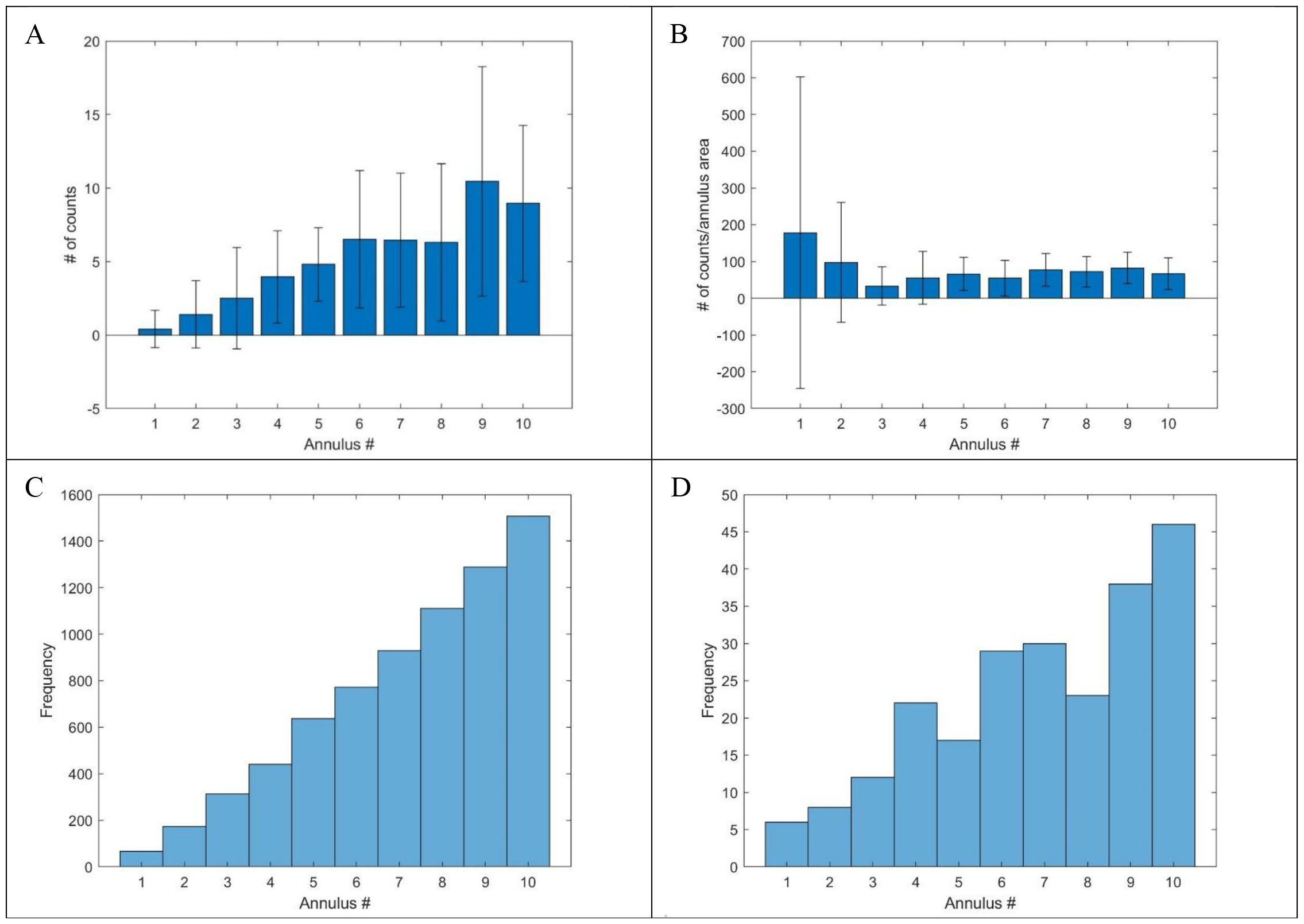
Localization of adhesive colloids in spinning suspension. (A) Positions and (B) normalized positions of colloids on annulus overlay (Fig. 1C) at end of simulation. Plotted are means ± standard deviations. N=20 for each data point. (C) Distribution of small size (≤7 colloids) particles and (D) distribution of large size (>7 colloids) particles on annulus overlay at end of simulation. Plotted are the sums of 500 simulation runs.

#### Case 3: Self-replicating adhesive colloids in spinning suspension

Particles were then allowed to replicate at every defined time interval. Replication was defined as a doubling of the number of particles in the simulation. Regardless of the size of the particle, the resultant would be the original particle and a particle of size 1 colloid. The number of particles decreased until a replication step, after which followed an increase in the number and size of particles with >1 colloid (Fig. 4A-H). All particles appeared uniformly distributed about the origin, and no angular bias was detected. Particle number decreased in a non-linear fashion with respect to time at each doubling (Fig. 5A). The average particle number approached an asymptote over an extended timeframe regardless of number of doublings (Fig. 5B). The most common particle size was 1 colloid; the second most common size of 2 colloids was ∼7X less frequent (Fig. 5C). Colloid number increased linearly with radial distance from origin, while colloid density remained uniform with all annuli (Fig. 5D,E). High particle density variability was again observed with the annulus closest to the origin. Small (≤7 colloids) and large (>7 colloids) sized particle frequencies also increased linearly with radial distance (Fig. 5F,G), indicative of uniform distribution.

**Figure 4:**
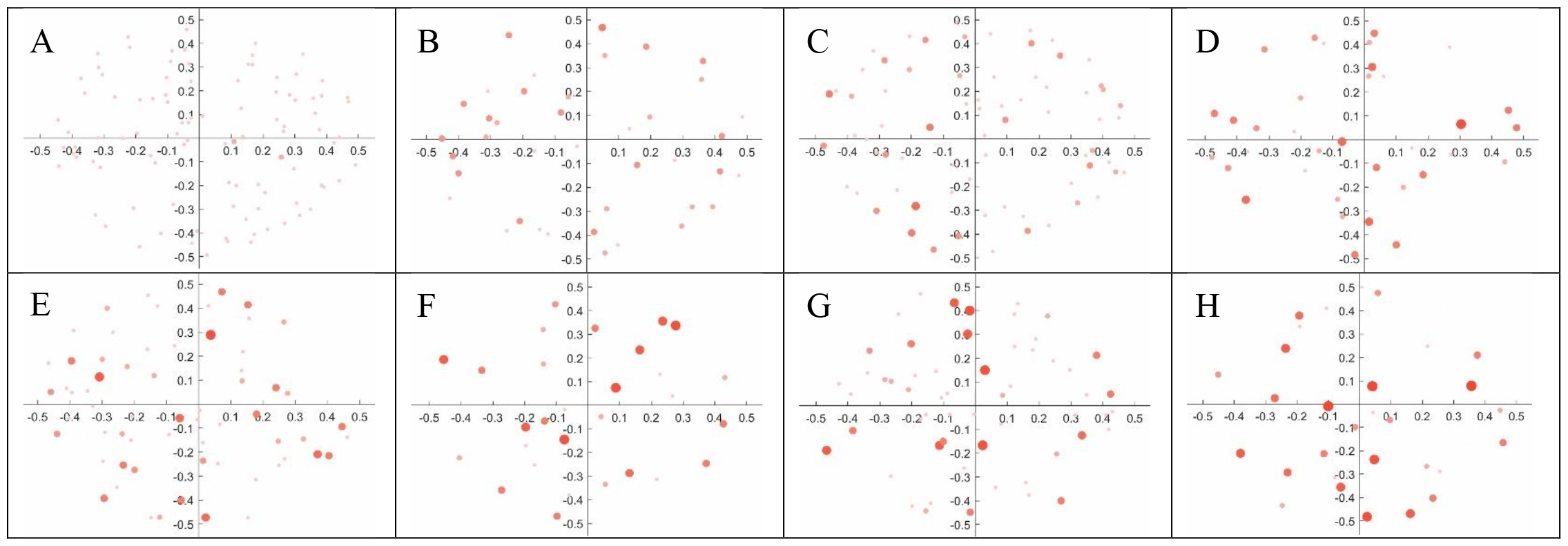
Self-replicating adhesive colloids in spinning suspension. Distribution of one hundred adhesive colloids at (A) the start, (B) before the first replication, (C) after the first replication, (D) before the second replication, (E) after the second replication, (F) before the third replication, (G) after the third replication, (H) end of the simulation. Size and color intensity correlate with number of adhered colloids.

**Figure 5:**
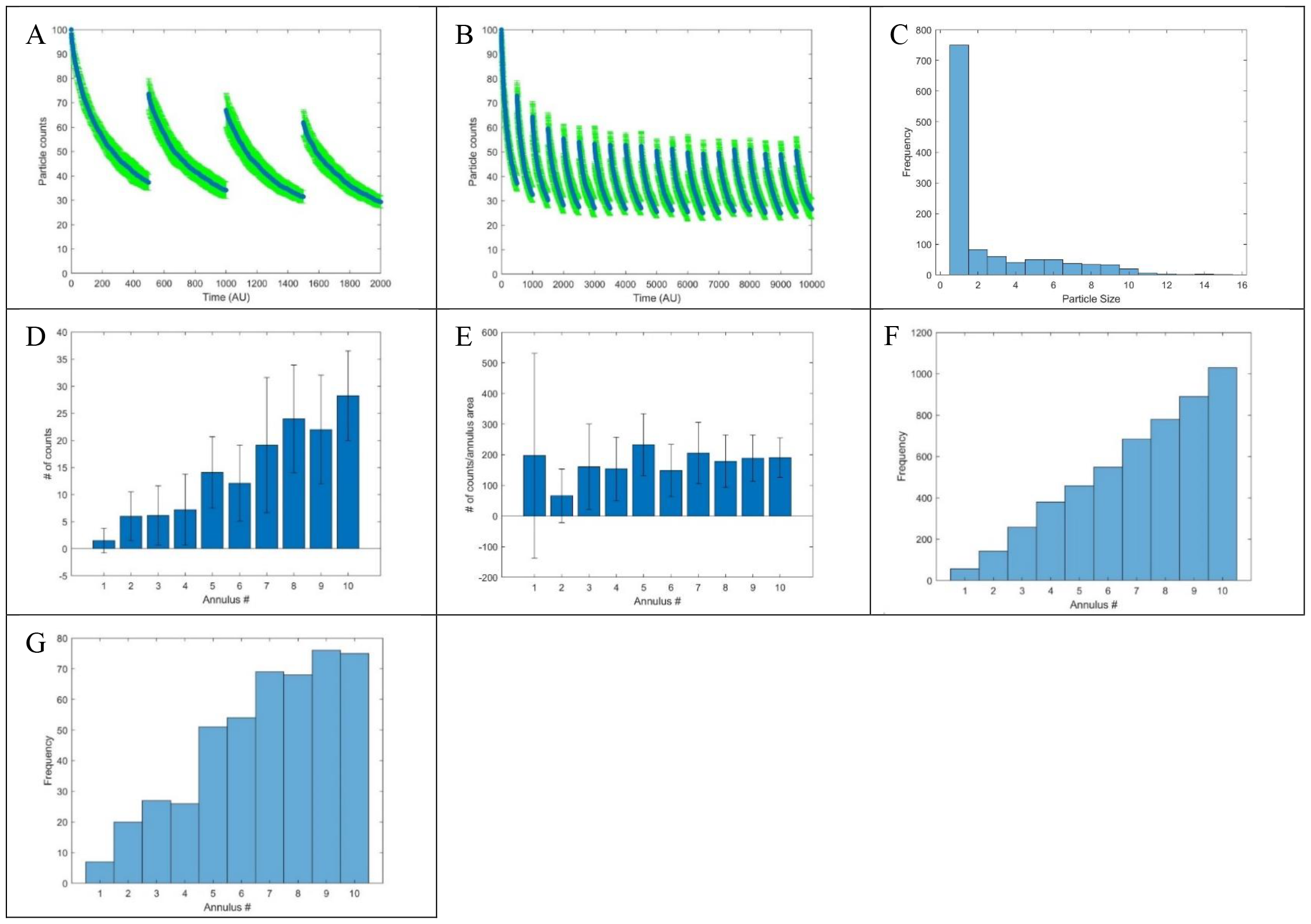
Localization of self-replicating adhesive colloids in spinning suspension. Change in particle count of self-replicating adhesive colloids in spinning suspension over (A) regular and (B) extended timeframes. Plotted are means ± standard deviations. N=20 for each data point. (C) Distribution of particle size at end of self-replicating colloid adhesion in spinning suspension simulation. Plotted is the sum of N=20 simulation runs. (D) Positions and (E) normalized positions of colloids on annulus overlay (Fig. 1C) at end of simulation. Plotted are means ± standard deviations. N=20 for each data point. (F) Distribution of small size (≤7 colloids) particles and (G) distribution of large size (>7 colloids) particles on annulus overlay at end of simulation. Plotted are the sums of 100 simulation runs.

## 3 Conclusion

Orbital shakers are one of the most common pieces of equipment found in any laboratory. The assumption behind their use is that continuous rotary shaking will quickly result in an equilibrium state of a homogenous mixture. Our simulation of colloid motion in an orbital shaker provided an estimate of the effect of orbital mixing over a short and long timeframe. Given different sets of common initial conditions and colloid interaction conditions, our simulation provided an estimate of colloid position and size distribution over time. We were able to visualize the dynamic equilibrium of particle count over time under conditions of particle self-replication and adhesion. Furthermore, our simulation setup was modular, thus allowing isolated testing of different initial and interaction conditions. This feature also allows for the substitution of different initial and colloid-colloid interaction requirements. Our assumptions for the ratios between colloid random walk speed, orbital shaker speed, minimum interparticle distance for adhesion, could be adjusted to fit different experimental setups. Initial colloid number and distribution could also be adjusted to simulate different experimental conditions. Adhesion and self-replication are also properties found in mammalian and bacterial cells. Their requirements could be adjusted to account for different biological phenomena; adhesion could be made dependent on particle size, different inter-particle distances, or a probability distribution based on custom variables relevant to the experiment. We present our MATLAB simulation as a scaffold on which these different adjustments can be supplemented to further researchers’ understandings of the internal dynamics of mixtures undergoing orbital shaking.

